# Systematic identification of secondary bile acid production genes in global microbiome

**DOI:** 10.1101/2024.06.08.598071

**Authors:** Yuwei Yang, Wenxing Gao, Ruixin Zhu, Liwen Tao, Wanning Chen, Xinyue Zhu, Mengping Shen, Tingjun Xu, Tingting Zhao, Xiaobai Zhang, Lixin Zhu, Na Jiao

## Abstract

Microbial metabolism of bile acids (BAs) is crucial for maintaining homeostasis in vertebrate hosts and environments. Although certain organisms involved in BA metabolism have been identified, a global, comprehensive elucidation of the microbes, metabolic enzymes, and BA remains incomplete. To bridge this gap, we employed hidden Markov models to systematically search in a large-scale and high-quality search library comprising 28,813 RefSeq multi-kingdom microbial complete genomes, which enabled us to construct a secondary bile acid (SBA)-production gene catalog. This catalog greatly expanded the distribution of SBA-production genes across 11 phyla, encompassing bacteria, archaea and fungi, and extended to 14 habitats spanning hosts and environmental contexts. Furthermore, we highlighted the associations between SBAs and gastrointestinal and hepatic disorders, including inflammatory bowel disease, colorectal cancer and nonalcoholic fatty liver disease, further elucidating disease-specific alterations in SBA-production genes. Additionally, we proposed the pig as a particularly suitable animal model for investigating SBA production in humans, given its closely aligned SBA-production gene composition. This gene catalog provides a comprehensive and reliable foundation on microbial BA metabolism for future studies, offering new insights into the microbial contributions to health and disease.

## Introduction

Microbial-derived secondary bile acids (SBAs) exert multifaceted influence on vertebrate hosts through various mechanisms. These mechanisms include direct cytotoxicity^1^, direct DNA damage^2^, and activation of receptors distributed across multiple tissues, including the liver, intestine, brain and breast^3–5^. SBAs originate from intestinal^6^ microbial enzymatic transformations on primary bile acids (PBAs) synthesized from cholesterol by host liver^7^. The principle microbial transformations of bile acids (BAs) comprise several key processes: deconjugation by bile salt hydrolases (BSHs)^8^, dehydroxylation by proteins encoded by bile acid-inducible (bai) genes^9–15^, oxidation and epimerization by position-specific hydroxysteroid dehydrogenases (α/β-HSDHs)^16^.

Disruption of BA microbial metabolism affects BA production and transport^17^, lipid and glucose metabolism^18^, as well as innate and adaptive immunity^19^, thereby contributing to the pathogenesis of a broad spectrum of diseases^20,21^. For instance, elevated deoxycholic acid (DCA) levels in the liver can facilitate hepatocellular carcinoma development by promoting the secretion of pro-inflammatory and tumor-promoting mediators^22^. In digestive tract, DCA also exacerbates intestinal inflammation by upregulating hepatic de novo BA synthesis^23^. Furthermore, certain animal-derived SBAs, such as hyodeoxycholic acid in pigs and ursodeoxycholic acid (UDCA) in bears, serve as therapeutic agents for conditions like nonalcoholic fatty liver disease (NAFLD)^24^ and fatal veno-occlusive disease^25^, respectively. Besides their important roles in the host, approximately 5% of BAs are released into the environment by vertebrate feces and urine^8,26^, serving as carbon– and energy-rich growth substrates for microorganisms in soil and water^27^. The isolation of environmental BA metabolizing microorganisms^28,29^ reveals the ubiquity of microbial BA metabolism and highlights how BAs and microorganisms from various hosts and environments can interconnect through food webs^30^. Microorganisms form complex ecological relationships^31^ and complement each other’s BA pathways, broadening the BA production repertoire^32^. Thus, both bottom-up control mediated by food and top-down control triggered by the impact of predation on prey in the ecosystem^33^, have the potential to influence the cycle and diversity of BAs.

Several microorganisms capable of BA metabolism have been reported, mainly including *Bifidobacterium*^34^, *Enterococcus*^35^ and *Listeria*^36^ for deconjugation; *Clostridium*^37^ for dehydroxylation; *Eggerthella*^38^ and *Ruminococcus*^16^ for oxidation and epimerization. Initially, the identification of BA metabolism enzymes in microorganisms relied primarily on biochemical analyses like enzyme activity assay^39^ and immunoblot analysis^40^, which were relatively inefficient. However, advancements in sequencing technologies and bioinformatics have revolutionized this field, facilitating the identification of specific BA metabolism processes within microbial genomes of single host gut through methods like BLAST and MUSCLE. For instance, BSH was identified in human fecal metagenomic datasets^41^, BaiE in whole-genome shotgun assembly sequences of human gut microbiomes^42^, and 7α/7β-HSDH in black bear fecal metagenomic datasets.^43^ Almut Heinken *et al.* expanded the scope of research by systematically identifying microbial BA deconjugation and biotransformation in the human gut with MUSCLE. However, their study only considered 693 human gut microbial genomes^32^, far fewer than the human gut microbiome reference set of 3,594 high-quality species genomes reported in 2022^44^. Despite these advances, most studies have focused on specific BA metabolism functions and limited sets of microbial genomes from single sources, which hinders our comprehensive understanding of the intricate global ecological network of microbial BA metabolism.

In response to these challenges, we aimed to expand the knowledge of BA microbial metabolism in the global microbiome. Utilizing a comprehensive database of 28,813 multi-kingdom microbial complete genomes from RefSeq database and employing hidden Markov models (HMMs), we constructed an SBA-production gene catalog. This catalog encompassed key genes from major pathways, including BSHs, Bai genes and HSDHs. This function-rich catalog, derived from a diverse array of cultured microorganisms not limited to specific habitats, can serve as a reliable reference for annotating SBA-production genes in global metagenomic sequencing data, and further assist us in systematically exploring the BA metabolism between species or communities.

## Results

### Global map of secondary bile acid-metabolic gene-microorganism

#### Overview of secondary bile acid production gene catalog

To construct a more comprehensive gene catalog associated with SBA metabolism, we generated HMMs for 13 gene families involved respectively, and performed a systematic search across 28,813 complete genomes of bacteria, archaea, and fungi in the RefSeq database. Rigorous screening criteria, based on the significance (e-value) and similarity (HMM score) of each hit (additional details are in Methods), ensured the reliability and accuracy of our findings. Ultimately, the SBA-production gene catalog contained a total of 1668 BSH genes, 241 Bai genes, 159 3αHSDH genes, 136 3βHSDH genes, 2770 7αHSDH genes, 7 7βHSDH genes, and 386 12αHSDH genes (Supplementary Figure 1, Figure 1a).

**Figure 1.**
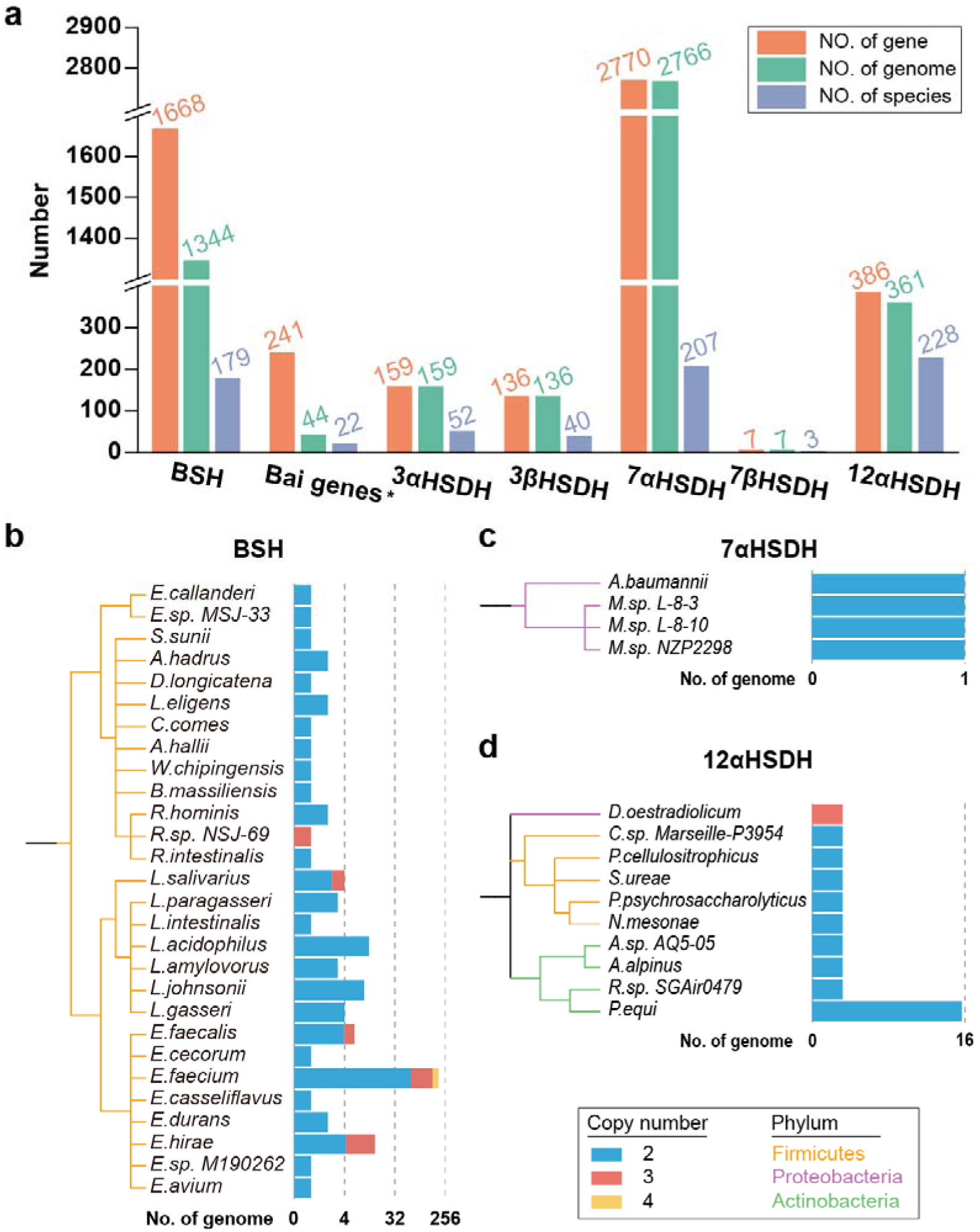
Overview of secondary bile acid production gene catalog. (**a**) The number of SBA-production genes (orange) as well as microbial genomes (green) and species (purple) carrying SBA-production genes. Phylogenetic trees of species with genome carrying duplicate (**b**) BSH, (**c**) 12αHSDH and (**d**) 7αHSDH genes. The branch colors represent different phyla. Stacked bar charts aligned to tree tips represent the number of genomes with duplicate genes. The blue, pink and yellow bar indicate the number of genomes with different copy number.

Further analysis revealed variability in the copy number of these genes (Figure 1a). Notably, 3αHSDH, 3βHSDH, and 7βHSDH were typically single-copy genes in these genomes. By contrast, BSH, 7αHSDH, and 12αHSDH exhibited multi-copy in certain genomes. BSH gene duplication was particularly prevalent, appearing in up to 271 genomes across 28 species of the Firmicutes phylum, with some genomes, such as eight from *Enterococcus faecium*, containing up to four copies. Additionally, multi-copy expression of BSH was relatively common in *E. faecium* genomes (Figure 1b), with 77.3% (194 of 251 genomes) exhibiting this trait. Regarding 7αHSDH, only two copies were identified within the Proteobacteria phylum, across one genome belonging to the *Acinetobacter* genus: GCF_020912005.1 (*Acinetobacter baumannii*) and three genomes from *Mesorhizobium* genus: GCF_013170825.1 (*Mesorhizobium sp. NZP2298*), GCF_016756635.1 (*Mesorhizobium sp. L-8-10*), and GCF_016756615.1 (*Mesorhizobium sp. L-8-3*) (Figure 1c). Notably, genomes with multi-copy 12αHSDH were distributed in three phyla: Actinobacteria, Firmicutes, and Proteobacteria. *Denitratisoma oestradiolicum* genome (GCF_902813185.1) within the Proteobacteria phylum had three copies, while the remaining 23 genomes had two (Figure 1d). *Prescottella equi*, a member of the Actinobacteria phylum, was the primary source of genomes with two copies of 12αHSDH. These findings shed light on differences in the copy number distribution of genes related to SBA production in microorganisms, which could enhance our understanding of the varied metabolic capabilities and ecological competitiveness of microorganisms involved in this process.

#### Taxonomic distribution of secondary bile acid production genes

The broad coverage of microbial genomes and metabolic gene types allowed us to investigate the distribution range of SBA-production genes and further gain insights into the BA metabolism capabilities of various microbial species and even different strains. Leveraging the lineage information from the RefSeq database, we explored the taxonomic distribution of SBA production genes across various ranks.

BSH genes were predominantly present in Firmicutes (1374 genes, 82.4%) and Actinobacteria (248 genes, 14.9%). At more specific taxonomic ranks, the majority of BSH genes within Firmicutes were distributed among genera, such as *Enterococcus* (655 genes, 39.3%), *Listeria* (281 genes, 16.8%), and *Lactiplantibacillus* (157 genes, 9.4%) (Figure 2a). Notably, among these, 273 genes were identified in 98.9% (273 genomes) of all *Listeria monocytogenes* genomes for research, and 156 genes were found in 87.6% (156 genomes) of all *Lactiplantibacillus plantarum* genomes (Supplementary Table 5). This indicated that these species generally possessed a single copy of BSH, not limited to specific strains. Additionally, our findings also expanded the distribution of BSH in archaea. Apart from the previously reported species *Methanobrevibacter smithii* and *Methanosphaera stadtmanae*^32^, BSH genes were also discovered in species such as *Methanobrevibacter millerae* and *Methanobrevibacter olleyae* of the Euryarchaeota phylum, as well as *Candidatus Methanomethylophilus alvus* of the Candidatus Thermoplasmatota phylum (Supplementary Table 2). Intrigued by the multi-kingdom distribution of BSH genes, we conducted a phylogenetic analysis revealing similarities between the BSH enzyme sequences in archaea Euryarchaeota and specific bacterial Firmicutes enzymes (Supplementary Figure 2a), suggesting potential horizontal gene transfer (HGT) event across the widespread distribution of BSH.

**Figure 2.**
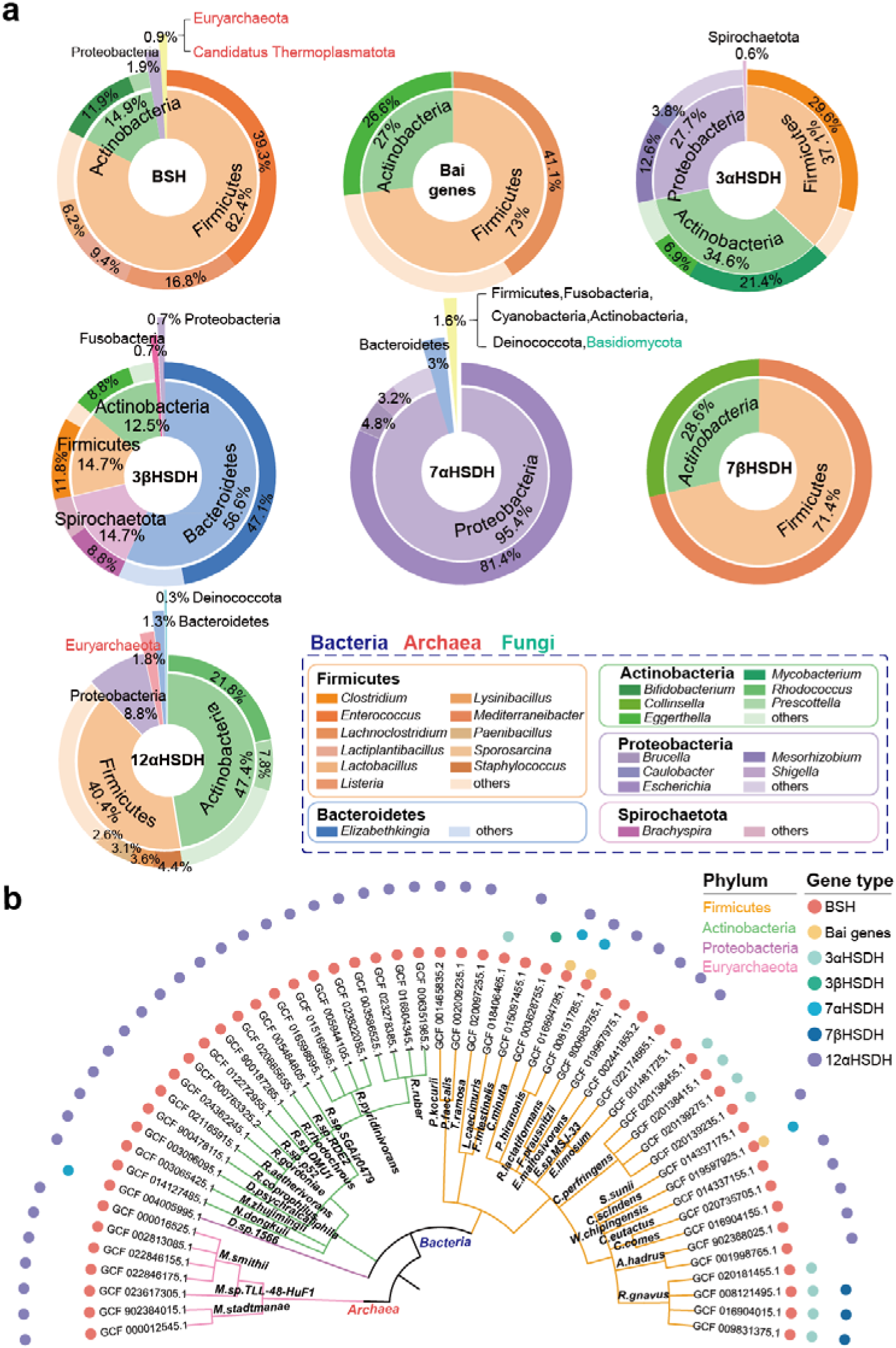
Taxonomy distribution of SBA-production genes. (**a**) Composition of different SBA-production genes. The pie charts show the proportions of genes across different phyla. The outer rings show the proportions of some main genera. (**b**) Phylogenetic tree of genomes carrying BSH and at least one Bai genes/HSDHs. The branch colors represent different phyla. Symbols aligned to tree tips represent different types of SBA-production genes.

Bai genes, previously reported to be sparsely distributed^45^, were found in only 22 species, originating from phylum Firmicutes (176 genes, 73.0%) and Actinobacteria (65 genes, 27.0%) (Figure 2a). Due to the cooperation nature of BA dehydroxylation involving multiple Bai genes, we further explored the taxonomic distribution of genomes encompassing all types of the Bai genes contained in our gene catalog simultaneously. We identified three genomes from the *Eggerthella* genus of the Actinobacteria phylum and 15 genomes primarily in the *Lachnoclostridium* genus (*Clostridium scindens*: 8 genomes, *Clostridium hylemonae*: 3 genomes) (Supplementary Table 2).

The taxonomic distribution of HSDHs varied depending on the subtype (Figure 2a). 3αHSDH predominantly existed in the *Clostridium* genus (47 genes, 29.6%) of Firmicutes phylum, the *Mycobacterium* genus (34 genes, 21.4%) of Actinobacteria phylum, and the *Mesorhizobium* genus (20 genes, 12.6%) of Proteobacteria phylum. Predominantly, *Clostridium perfringens* and *Mycobacterium avium* were the major carriers, with 88.5% and 97.1% of their genomes containing this gene, respectively (Supplementary Table 5). In contrast, 3βHSDH genes demonstrated a broader and distinctive distribution, predominantly within phylum Bacteroidetes (77 genes, 56.6%) and Spirochaetota (20 genes, 14.7%). Notably, *Elizabethkingia anophelis* emerged as a significant carrier, all 50 genomes of whom produced 3βHSDH (Supplementary Table 5), which suggested a core metabolic function of this species. 7αHSDHs displayed extensive distribution across seven bacterial phyla, with a staggering 95.4% (2642 genes) in Proteobacteria phylum, mainly concentrated in the species *Escherichia coli*, which accounted for 97.9% of the cases (2203 genomes) in our study (Supplementary Table 5). Interestingly, this gene was also found in the fungus *Rhizoctonia solani* of Basidiomycola phylum, which displayed phylogenetic similarity with enzymes in Proteobacteria (Supplementary Figure 2b), indicating a possible origin of fungi 7αHSDH. Conforming to previous research^45^, 7βHSDH enzymes were less prevalent, found primarily in *Ruminococcus gnavus* (4 genes) and *Ruminococcus torques* (1 gene) from Firmicutes phylum, and *Collinsella aerofaciens* (2 genes) from Actinobacteria phylum. In addition, 12αHSDH was predominantly synthesized by species within phylum Actinobacteria (183 genes, 47.4%) and Firmicutes (156 genes, 40.4%), with genes mainly distributed in the genus *Rhodococcus* (84 genes, 21.8%). There was also a sparse distribution of 12αHSDH in three species of Euryarchaeota archaea: *Methanobrevibacter smithii* (4 genes), *Methanobrevibacter sp. TLL-48-HuF1* (1 gene), and *Methanosphaera stadtmanae* (2 genes). In line with the potential HGT of BSH, these enzyme sequences in Euryarchaeota were also phylogenetically close to those of Firmicutes (Supplementary Figure 2c).

Our study uncovered 55 genomes from 37 species exhibited the simultaneous possession of the BSH and either the Bai genes or HSDHs, indicative of a more independent capability for SBA production. These genomes were primarily distributed among phylum Firmicutes (28 genomes, 50.9%), Actinobacteria (19 genomes, 34.5%) in bacteria, and Euryarchaeota in archaea (7 genomes, 12.7%). Among these, species like *Ruminococcus gnavus*, *Clostridium perfringens* (Firmicutes), *Rhodococcus pyridinivorans*, *Rhodococcus ruber* (Actinobacteria), and *Methanobrevibacter smithii* (Euryarchaeota), displayed up to four genomes capable of multifunctional SBA metabolism. Notably, *Devosia sp. 1566*. was the only multi-functional genome identified in the Proteobacteria phylum. From the perspective of the diversity of SBA metabolism enzymes, multi-functional microorganisms demonstrated the tendency of simultaneously possessing both BSH and 12αHSDH synthesis capabilities, with 43 genomes in 31 species displaying this trait. Moreover, genomes of Firmicutes showcased a more diverse range of SBA metabolism enzymes. Apart from the BSH-7αHSDH and BSH-12αHSDH combinations shared with other genomes, the Firmicutes genomes possessed six additional combinations involving other genes (Figure 2b). These functionally-rich genomes may play a significant role in the process of SBA metabolism and provide new insights for future research on biotechnological production of SBA using these microorganisms.

Altogether, our findings demonstrated that microorganisms with BA metabolism capabilities were widely distributed across different kingdoms, spanning bacteria, archaea, and fungi. However, the functions and distributions of these metabolic genes varied significantly, underscoring the importance of considering taxonomic characteristics when studying microbial bile acid metabolism.

#### Mammalian and environmental microbiota both served as reservoirs for secondary bile acid production genes

In order to obtain a comprehensive understanding of the distribution of genes involved in SBA metabolism across various hosts and environments, we conducted a thorough analysis using the Global Microbial Gene Catalogue (GMGC), a global-scale gene catalog constructed from worldwide metagenomes covering 14 habitats^46^. By leveraging this resource, we examined the habitats and geographical locations of these genes to provide a global perspective on their distribution.

The findings underscored the widespread distribution of SBA-production genes across a variety of global habitats. Notably, these genes were present not only in mammal hosts like humans, pigs, and mice, but also in diverse environmental settings such as wastewater, marine, and soil (Figure 3a). A comparative analysis of the proportions of these SBA-production genes in different habitats revealed intriguing variations in both prevalence and composition (Figure 3b). For instance, the gut microbiome of dogs exhibited the highest proportion of SBA-production genes, accounting for 0.02158% (793 genes), while marine displayed the lowest, as just 0.00003% (27 genes). Within mammal hosts, SBA-production gene prevalence varied notably among organs, reflecting the primary sites of bile acid metabolism. The gut displayed significantly higher proportions compared to less metabolically active sites such as the oral and nasal cavities. Specifically, the human gut contained 0.01126% (5,914 genes) of the total gene set, whereas the human oral cavity had only 0.00152% (203 genes). The environmental proportion of SBA-production genes appeared to be influenced significantly by the human activities. Wastewater environments showed a higher proportion of these genes (404 genes, 0.01471%) compared to built environments (281 genes, 0.00346%), freshwater (10 genes, 0.00036%), soil (230 genes, 0.00030%), and marine settings (27 genes, 0.00003%). This suggested a potential increase in microbial BA metabolism capacity where human impact is prevalent. Additionally, we observed differences in the proportional composition of specific bile acid metabolism genes, such as BSH, Bai genes, and HSDHs across these habitats. In guts of humans and pigs, BSH genes were more abundant, while HSDH genes were more prevalent in other environments.

**Figure 3.**
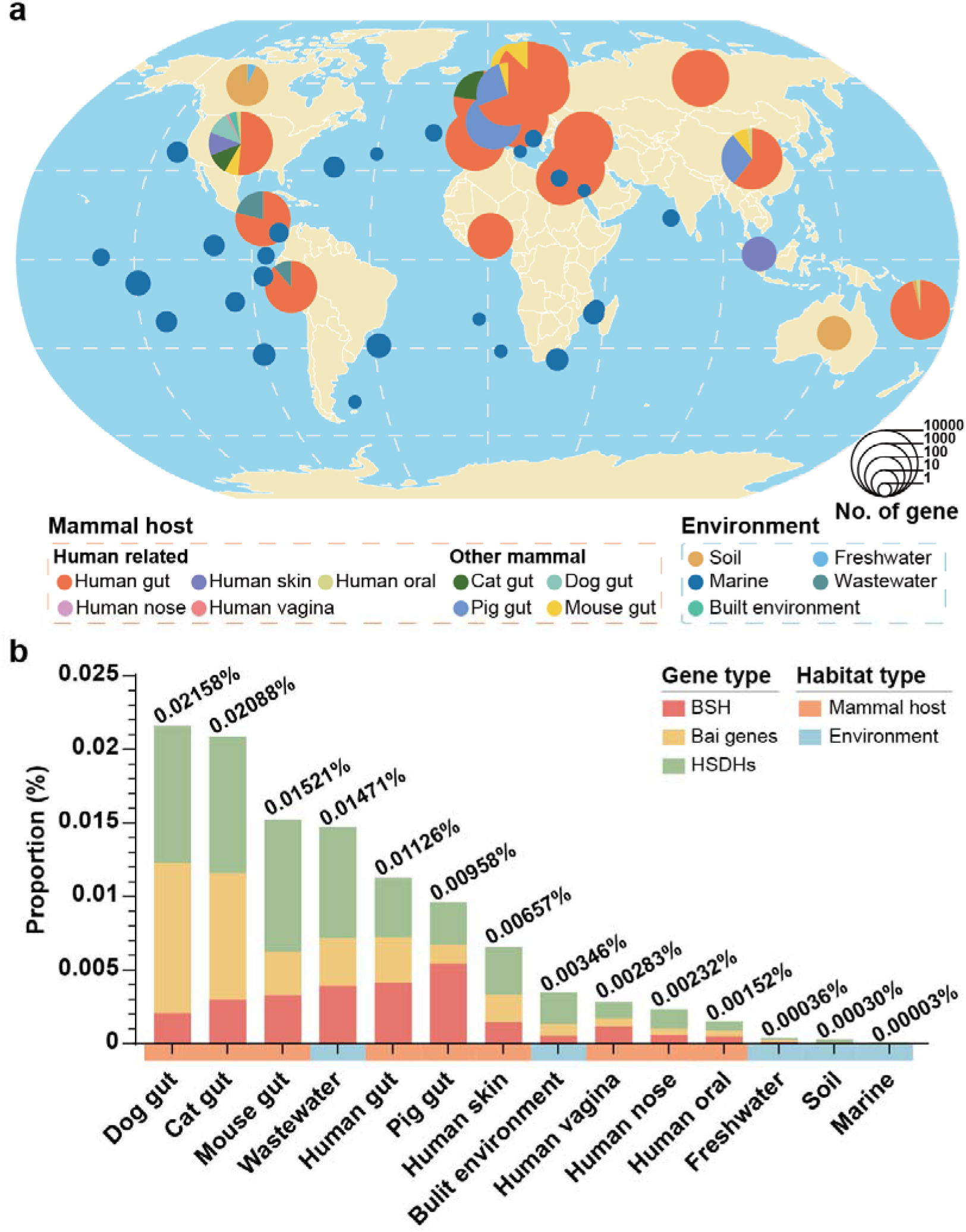
Habitat distribution of SBA-production genes. (**a**) Global map representing SBA-production genes in GMGC. The size of the pie chart represents the number of SBA-production unigenes. Different colors represent different habitats. (**b**) SBA-production genes composition in GMGC. Stacked bar chart shows proportions of BSH (red), Bai genes (yellow) and HSDHs (green) in total genes of different habitats.

This investigation highlighted that microbial BA metabolism was not restricted to specific organisms or ecosystems but rather a ubiquitous process across diverse hosts and environments. The variations in gene prevalence and composition likely reflected the adaptation of microbial communities to the available substrates or favored metabolic products in each habitat.

### Disease-specific alterations in the composition of secondary bile acid production genes

Considering the important role of BA metabolism in gastrointestinal and hepatic pathophysiology, we comprehensively elucidated the profiles of SBA production among inflammatory bowel disease (IBD), colorectal cancer (CRC) and NAFLD based on gut microbial genes and species.

We compared the overall disruption levels of SBA metabolism across different diseases utilizing the defined SBA metabolism differential score, which incorporated the proportion of each gene type and the significance of their differences between disease and control group. In general, disruptions of SBA metabolism showed significant alterations in intestinal diseases such as IBD and CRC, particularly in Crohn’s disease (CD) and CRC, followed by ulcerative colitis (UC), while adenomas exhibited relatively smaller changes. Moreover, due to the gut-liver axis, NAFLD was also found to be associated with changes in SBA metabolism (Figure 4).

**Figure 4.**
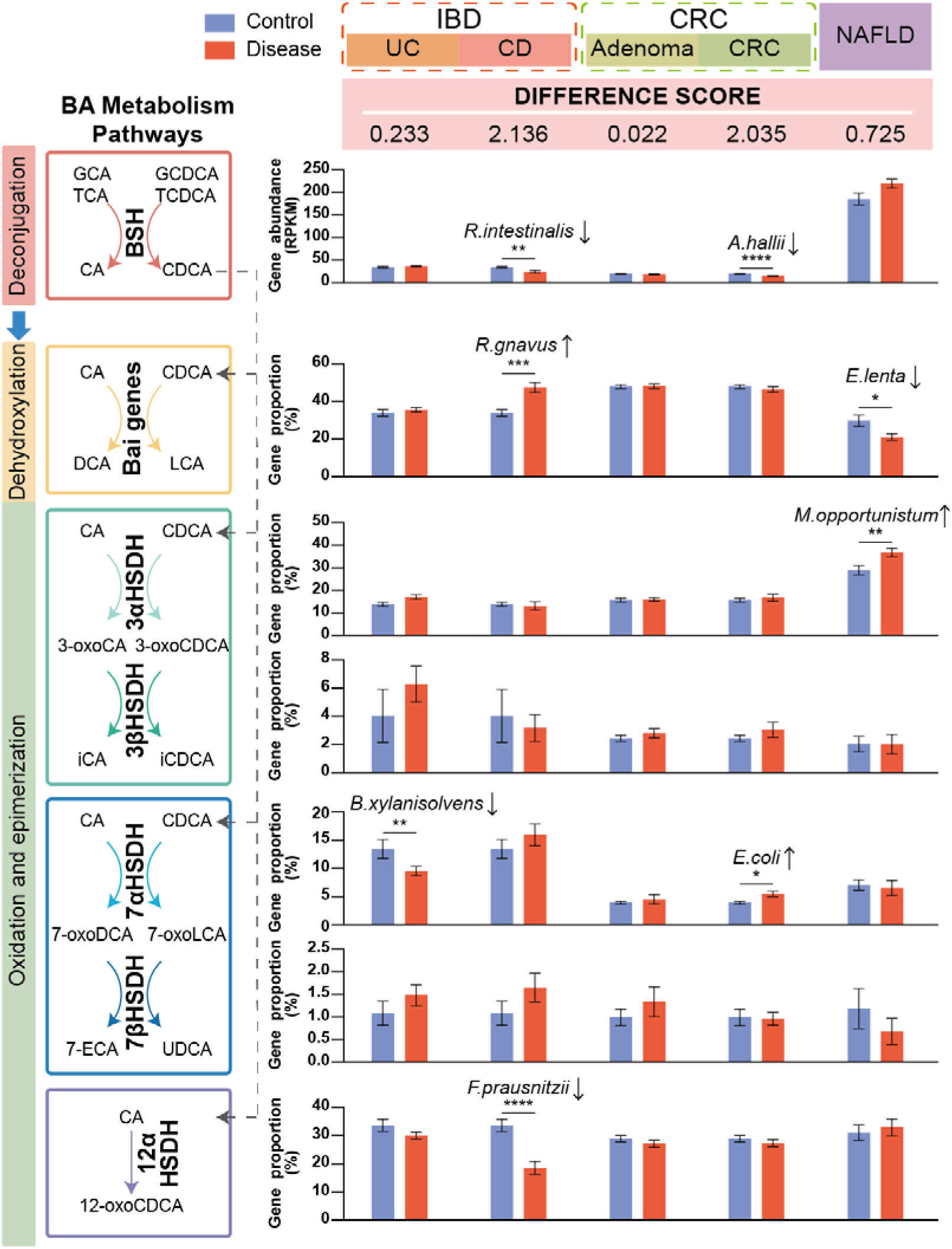
Profiles of SBA-production genes in intestinal and liver diseases. The bar plots show the gut microbial gene abundance of BSH as well as proportions of Bai genes and HSDHs in different disease states. The microorganisms located above the bar plot are the major differential species possessing this gene and are consistent with the gene alteration. The definition of ‘Difference score’ is in equation (3). Data are shown as mean with SE. The statistical differences between groups were determined by two-tailed Mann-Whitney U-test (UC, CD, adenoma, CRC) or paired t test (NAFLD), the p values were converted to asterisks(* for p < 0.05; ** for p < 0.01; *** for p < 0.001 and **** for p < 0.0001). Number of samples by disease state for each study were: IBD: control = 23, UC = 124, CD = 21; CRC: control = 63, adenoma = 47, CRC = 46; NAFLD: control = 10, NAFLD = 10.

We further detailed the distinct differences in metabolic gene profiles across various diseases, providing insights into how each condition uniquely influences SBA metabolism (Figure 4). BSH genes serve as gateways for SBA metabolism, and their gene abundances were significantly downregulated in patients with CD and CRC. Notable BSH-possessing microorganisms such as *Roseburia intestinalis*, *Anaerobutyricum hallii*, *Blautia sp. SC05B48* showed a consistent decrease(Supplementary Table 8). In CD, *R.intestinalis* exhibited a higher relative abundance weighted by copy number (Supplementary Figure 3b), while in CRC it was *A.hallii* (Supplementary Figure 3e).

Next, we considered the subsequent biotransformation of dehydroxylation, oxidation and epimerization as a whole, and compared the proportional composition of the Bai genes and various HSDH genes. Notably, we observed significant proportion changes specific to different diseases. The proportion of Bai genes, crucial for the generation of DCA and LCA through dehydroxylation, showed significant variation. In comparison to the control group, it significantly increased in CD, whereas notably decreased in NAFLD. *Ruminococcus gnavus* was predominantly increased in CD (Supplementary Figure 3c), while *Eggerthella lenta* showed a significant decrease in NAFLD (Supplementary Figure 3g). α-HSDHs carry out the oxidation of the hydroxyl group at the 3-, 7-, and 12-carbons of cholic acid (CA) or chenodeoxycholic acid (CDCA), a significant increase in 3αHSDH was observed in NAFLD, notably consistent with the alteration in weighted relative abundance of *Mesorhizobium opportunistum* (Supplementary Figure 3h). The 7αHSDH, significantly decreased in UC, primarily affected by the producer *Bacteroides xylanisolvens* (Supplementary Figure 3a), but in CRC with a rise in the proportion of this gene, aligning with a significant increase of the high-abundance 7αHSDH-possessing species *Escherichia coli* (Supplementary Figure 3f). The 12αHSDH, exclusively altered in CD, showed a decrease correlating with a drop in the weighted relative abundance of *Faecalibacterium prausnitzii* producing 12αHSDH (Supplementary Figure 3d). Additionally, no significant changes were observed in these diseases regarding the proportions of 3βHSDH and 7βHSDH, which catalyze reduction after the oxidation by α-HSDH.

We further explored the species composition of different genes based on the weighted relative abundance of species possessing SBA-production genes (Supplementary Figure 4). Species of Firmicutes phylum emerged as the primary SBA producers in the gut, with the highest proportion in BSH, Bai genes, and 12αHSDH. Furthermore, Actinobacteria primarily synthesized 3αHSDH, while Bacteroidetes were the main source of 3βHSDH and 7αHSDH genes. Notably, *Ruminococcus gnavus*, *Ruminococcus torques*, and *Collinsella aerofaciens* with 7βHSDH synthesis ability were found in the gut. Additionally, the archaea producing BSH and 12αHSDH, as well as fungus producing 7αHSDH identified in the gut also play roles in BA metabolism, contributing to the diversity and functionality of the microbial community.

Taken together, we systematically elucidated the profiles of SBA production in different diseases from the perspectives of both metabolic genes and species, and further identified disease-specific alterations in SBA-production genes. The findings emphasized the important role of SBAs and their microbial metabolism in gastrointestinal and hepatic diseases.

### The metabolic process of secondary bile acids in pigs was more similar to that in humans

Animal models are critical in human medical research. To evaluate the feasibility of utilizing animal models for studying human SBA metabolism, we devised a score that quantified the resemblance between the intestinal microbial SBA production genes of various animals and humans. This score reflected not only the direct match of key SBA metabolism genes but also their relative contributions to the overall metabolic process.

Among the evaluated animal models, pigs displayed the greatest overall similarity to humans in terms of SBA metabolism, followed by cats, while mice and dogs exhibited lower similarity scores (Figure 5). Specifically, the resemblance between pigs and humans in terms of SBA-production genes in the intestines was primarily observed in BSH, 3αHSDH, 7βHSDH, and 12αHSDH, with 764, 200, 24, and 132 same genes respectively, which represented 35.2%, 31.7%, 49.0%, and 13.3% of the corresponding genes in humans. Conversely, the number of unique genes in humans was relatively small, including only 146 (6.7%) BSH, 32 (5.1%) 3αHSDH, 1 (2.0%) 7βHSDH, and 64 (6.4%) 12αHSDH. However, cats’ microbiome showed higher similarity scores for Bai genes and 7αHSDH. Of the 1627 Bai genes in humans, 302 were identical to those found in cats, and 1192 showed similarity. For the 304 7αHSDH genes, 51 were identical and 206 were similar. It was worth noting that the similarity of 3βHSDH between the animal models and humans was low. Only 56 genes were shared across the four animal models (41 in pigs, 33 in cats, 3 in mice, and 32 in dogs), and 32.6% (46 genes) of human intestinal 3βHSDH was unique.

**Figure 5.**
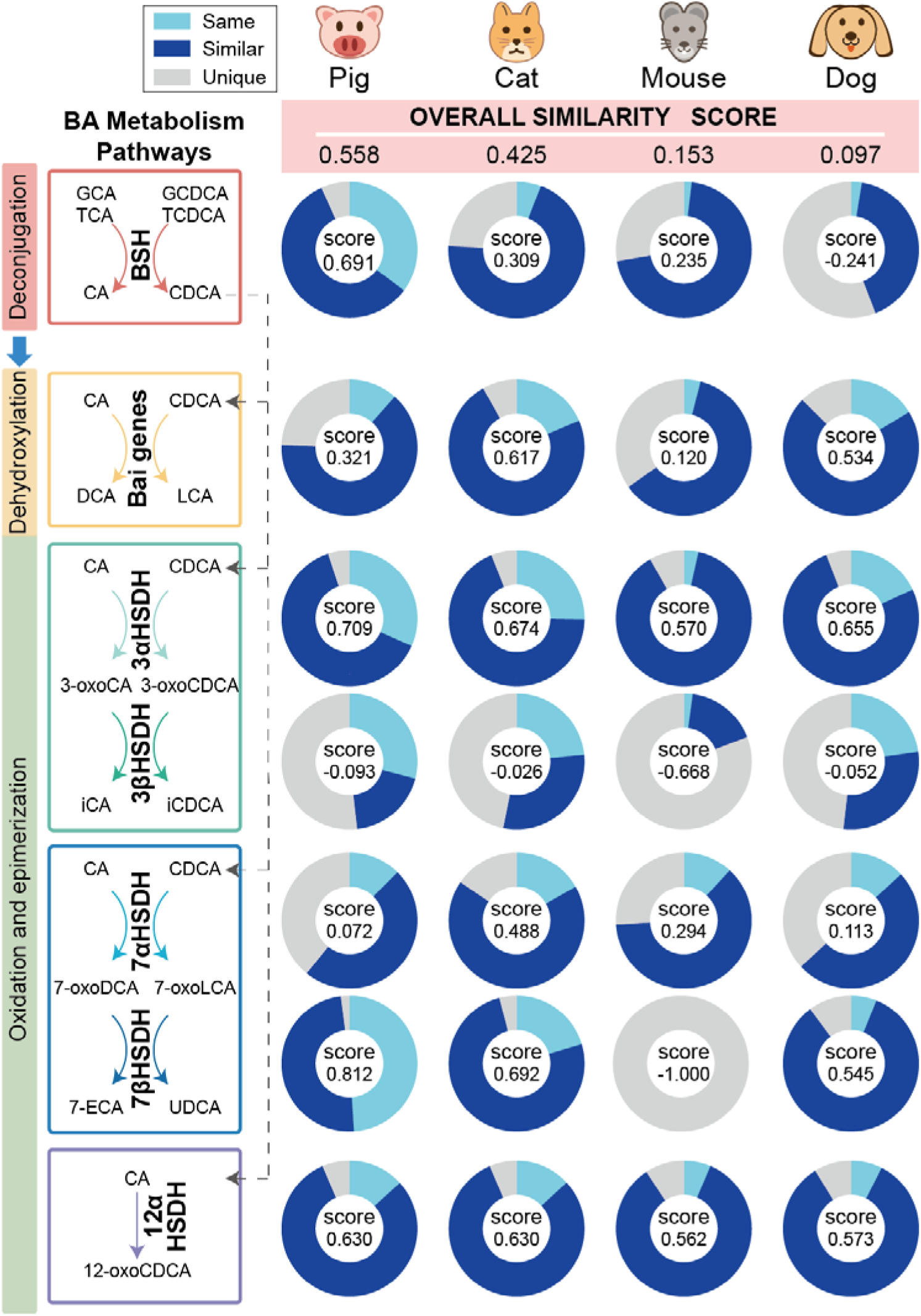
Comparison of SBA-production genes between humans and different animal models. The pie charts show the proportions of different kinds of SBA-production genes in total human genes across four animal models. ‘score’ in pie charts is the ‘Single similarity score’ defined in equation (4). The definition of three kinds of genes is in Methods, and ‘Overall similarity score’ in equation (5).

These findings underscored the variable degree of similarity in SBA-production genes between human and common animal models. Pigs exhibiting the highest similarity score, emerged as particularly suitable for modeling human SBA metabolism. This also highlighted the importance of selecting animal models that closely mimic human metabolic processes for other studies, which is crucial for the translational relevance of biomedical research.

## Discussion

Accurate and comprehensive gene definition is essential for the analysis of functional and ecological roles of microbial communities. In this study, we systematically identified genes involved in BA deconjugation, dehydroxylation, oxidation, and epimerization among 28,813 complete multi-kingdom microbial genomes sourced from the RefSeq database. Our expanded SBA-production gene catalog builds upon and refines previous studies^8–16,32,34–41,43,47–51^, filling the gap in limited microbial and enzymatic resources. This enhancement allows for a more thorough exploration of the SBA metabolism from both microbial species and functional perspectives across diverse habitats, enriching our understanding of the ecological impact of these processes globally.

Our gene catalog revealed a broad taxonomic distribution of SBA-production genes across multi-kingdom microorganisms, with a distinct tendency for microorganisms to participate in specific stages of SBA metabolism. This suggests that effective SBA metabolism often requires inter-microbial cooperation. Notably, 55 genomes from 37 species demonstrated multifunctionality in SBA metabolism, with *Peptacetobacter hiranonis* from the Firmicutes phylum exhibiting the most enriched functionality. It simultaneously possessed the synthesis potential of BSH, Bai genes, 7αHSDH, and 12αHSDH. This species, already identified as a biomarker for gastrointestinal functionality in dogs^52^, and showed strong and significant correlations with BaiCD, and the relative concentration of SBA in dogs^53^. Future research can concentrate on these multifunctional SBA-production microorganisms, as they hold potential for deeper investigation due to their central role within microbial communities.

The varying frequency and composition of SBA-production genes across different habitats, underscored the critical interplay between microbial communities, host physiology, and environmental factors. The widespread distribution of SBA-production genes across various habitats, notably in mammalian hosts and human-impacted environments, highlights the influence of environmental factors on microbial BA metabolism patterns, which may, in turn, impact disease pathogenesis. Mass spectrometry reanalyses^21^ have detected microbe-derived BAs in organs beyond the liver and intestines, such as the skin^54^, suggesting broader systemic roles. Therefore, the comparative analysis of SBA-production genes in different organs can facilitate a better understanding of the diverse physiological functions of BAs. Microbial BA metabolism in the environment can produce hormone-like metabolites^27^, suggesting that variations in SBA-production genes composition may have ecological effects.

With the comprehensive knowledge of the species and genes involved in BA metabolism, we could explore the related issues from a fresh perspective. One of the applications was to systematically analyze the alterations in microbial BA metabolism that occur under various conditions like diseases. In our study involving multiple gastrointestinal and hepatic disease cohorts from Europe, we specifically analyzed the profiles of SBA-production genes to investigate their role in various diseases. Our findings uncovered disease-specific changes in SBA metabolism profiles. For patients with IBD, the capability of their gut microbiota in synthesizing hydrophilic and less toxic SBAs through α-HSDHs was weakened in both UC and CD. However, distinct differences were found between these two subtypes, with CD showing a greater overall change. Specifically, CD was characterized by a substantial decrease in the BSH gene abundance and an increase in the Bai genes proportion. This pattern is consistent with previous findings of reduced SBAs^55,56^ and elevated Bai genes abundance^57^ in CD, supporting theories of inflammation-associated and CD-specific metabolic disruptions^55^. These insights enhance our understanding of IBD pathogenesis and may inform more targeted research into the metabolic underpinnings of UC and CD. Furthermore, our study highlighted the importance of SBA metabolism in the progression from health to adenoma and ultimately to CRC. While adenomas did not show significant changes, CRC cases exhibited a reduced capacity to synthesize SBAs, yet an increased proportion of hydrophilic SBA like UDCA produced by 7α/βHSDH. This may be linked to elevated supraphysiological hydrophilicity levels and potentially increasing the risk of developing CRC^58^. In contrast to gastrointestinal diseases, NAFLD exhibited an increase in BSH genes, although this change was not statistically significant. Microbe-derived BAs weakly induced FXR activation and notably reduced CDCA-induced FXR activation^59^. Thus, the result above corresponded to our previous study that highlighted the suppression of FXR-mediated signaling in NAFLD^20^. Additionally, we noted a trend within the NAFLD gut microbiome toward producing more SBAs with enhanced hydrophilicity, driven by significant alterations in the proportion of Bai genes and 3αHSDH. Overall, alterations in SBA-production genes can potentially impact disease development and progression by altering the abundance of SBAs and their overall hydrophobicity, toxicity, and receptors interactions. This highlight the potential of SBAs and their metabolic pathways as targets for therapeutic intervention. Beyond merely examining the overall content of SBAs, it is crucial to explore the compositional changes in different types of SBAs to understand their roles in various diseases fully. Analyses based on fecal metagenomic sequencing provide insights into these changes but only offer a partial view. Therefore, the establishment of additional metabolomics cohorts is essential for a more comprehensive understanding of how disruptions in SBA metabolism affect different diseases.

Our gene catalog could also facilitate the comparison of SBA-production genes across different habitats. This comparative study, particularly among common animal models, namely pig, mouse, cat, and dog, assessed their suitability for BA research. Pigs were found to be the more suitable animal model due to their close resemblance to human microbial BA profiles. The microbial composition of pig feces shares similarities with human^60^, with 96% of the functional pathways identified in the human gut microbiome gene reference catalog are present in the pig catalog^61^. Moreover, pigs exhibit comparable physiological characteristics in gastrointestinal tract development and digestive function^62^, mirroring human dietary, digestive patterns^63^. These findings support the adoption of pigs as a valuable model for human biological and disease study. Future research should expand the comparative scope to more animals like bears, addressing our limitation in using GMGC. Additionally, considering variations in types and proportions of PBAs among different mammalian species^64^ could fully capture the complexity of BA metabolism.

While our catalog significantly advances the field, it does not encompass the complete diversity of microbial-derived BAs recently revealed^65^. Emerging findings have discovered novel functionalities of known enzymes involved in BA transformations, such as BSH for conjugating various amino acids to BAs^66^, as well as the discovery of previously unknown enzymes like BAS-suc, responsible for producing 3-succinylated cholic acid^67^, which suggests a vast uncharted territory of microbial metabolism. Future efforts should aim to uncover these new metabolic pathways and integrate them into our existing frameworks to provide a more comprehensive view of microbial contributions to health and disease.

In conclusion, this systematic gene catalog derived from abundant microbial genomes highly enriches microbial SBA production resources and further promotes the identification of related genes in the global microbiome. providing valuable context for further research in this field. By applying our findings to various disease contexts and animal models, we offer new perspectives on the potential roles of microbial metabolism in health and disease, paving the way for innovative therapeutic approaches based on microbial and metabolic manipulation.

## Methods

### Constructing the secondary bile acid production gene catalog

To better identify the target genes, HMM, an exhaustive algorithm based on dynamic programming^68^, was used for its greater sensitivity compared to heuristic algorithms and capability to capture position-specific alignment information^69^. HMM seed protein sequences of BSH, Bai genes and HSDHs derived from different species (Supplementary Table 1) were obtained from the PubSEED database^70,71^, and then aligned in Clustal Omega v1.2.4^72^ to construct HMMs on full-length proteins via hmmbuild (default mode, HMMER 3.3.2, hmmer.org) respectively. These model seed sequences were realigned to the models using hmmalign (default mode) before rebuilding models based on the alignments, and this iterative process was repeated three times to ensure the robustness of models. Subsequently, these HMMs were used to search for orthologs (hmmsearch (–tblout)) in the protein sequences of all 28,813 complete microbial genomes (bacteria: 28,324 genomes; archaea: 466 genomes; fungi: 23 genomes) provided by the National Center for Biotechnology Information Refseq database (accessed in July 2022). Hits with e-value less than 1e^−65^ were selected as candidates. To ensure high-quality sequences, further sorting was done based on HMM score from hmmsearch results, and cutoffs (BSH: 400; BaiA: 390; BaiB: 1000; BaiCD: 1090; BaiE: 360; BaiF: 1000; BaiG: 950; BaiH: 1400; 3αHSDH: 300; 3βHSDH: 340; 7αHSDH: 325; 7βHSDH: 580; 12αHSDH: 250) at obvious HMM score drops were set to define secondary bile acid production genes. For Bai genes and 7βHSDH, due to limited high-score results, genes from the PubSEED database were included as part of the secondary bile acid production gene catalog (Supplementary Table 2). The DNA sequences corresponding to the SBA-production gene catalog were in Supplementary Table 3. Due to the redundancy in the protein sequences encoded by these genes, we constructed a non-redundant protein catalog based on their protein accession numbers (BSH: 416 sequences; Bai genes: 146 sequences; 3αHSDH: 70 sequences; 3βHSDH: 62 sequences; 7αHSDH: 319 sequences; 7βHSDH: 3 sequences; 12αHSDH: 265 sequences, Supplementary Table 4). This non-redundant protein catalog served as comprehensive and high-quality reference sequences for subsequent analysis of SBA-production microbial enzymes in metagenomic data.

### Phylogenetic analyses of BSH, 7αHSDH and 12αHSDH genes

To understand the evolution of BSH, 7αHSDH and 12αHSDH distributed across different microbial kingdoms, multiple sequence alignments were performed by Clustal Omega v1.2.4 with non-redundant protein sequences of BSH, 7αHSDH and 12αHSDH and trimmed with trimAl v1.4^73^ on automated1 mode. Next, maximum-likelihood phylogenetic trees were inferred by IQ-TREE v2.2.6^74^ using the suggested best-fit model (BSH: LG + R6; 7αHSDH: Q.pfam + R6; 12αHSDH: Q.pfam + R7) with 1000 ultrafast bootstrap replicates and visualized and annotated using iTOL^75^.

### Analyzing the habitat distribution of secondary bile acid production gene catalog

For the distribution of SBA-production genes at the global scale, DNA sequences of 302,655,267 unigenes from GMGC v1^46^ were used as the query for BLASTX^76^ searches against the non-redundant protein catalog of SBA-production. Only blast hits with at least 70% identity and less than 1e^−50^ e-value were considered quality hits (Supplementary Table 6). By combining the habitat and geographic location information provided by the GMGC database for the sources of these unigenes, we analyzed the habitat and geographic distribution of genes involved in SBA metabolism.

### Metagenomic data processing

Raw sequencing data were downloaded from the Sequence Read Archive (SRA) using the following identifiers: PRJEB1220 (IBD) for Nielsen et al.^77^, ERP008729 (CRC) for Feng et al.^78^ and PRJNA420817(NAFLD) for Mardinoglu et al.^79^ (in this study, paired ‘NAFLD’ and ‘Control’ samples were obtained from fecal samples collected on day 0 and day 14 of participants followed an isocaloric low-carbohydrate diet with increased protein content for 14 days). Firstly, KneadData v.0.6 was utilized to remove low-quality and contaminant reads including host-associated ((hg38, felCat8, canFam3, mm10, rn6, susScr3, galGal4 and bosTau8 from UCSC Genome Browser) and laboratory-associated sequences by Trimmomatic v0.39^80^ (SLIDINGWINDOW:4:20 MINLEN:50 LEADING:3 TRAILING:3) and bowtie2 v.2.3.5^81^. Thereafter, filtered reads were used to generate taxonomic and functional profiles. Taxonomic classification of bacteria, archaea, fungi and viruses was performed against our pre-built reference database using Kraken2^82^. The pre-built database comprises 32,875 bacterial, 489 archaeal, 11,694 viral reference genomes from the National Center for Biotechnology Information (NCBI) RefSeq database (accessed in August 2022), and 1,256 fungal reference genomes from the National Center for Biotechnology Information Refseq database, FungiDB (http://fungidb.org) and Ensembl (http://fungi.ensembl.org) (accessed in August 2022). It was built using the Jellyfish program by counting distinct 31-mer in the reference libraries, with each k-mer in a read mapped to the lowest common ancestor of all reference genomes with exact k-mer matches. And we used Bracken v.2.5.0^83^ to accurately count taxonomic abundance. The read counts of species were converted into relative abundance for further analysis. For function profiles, reads were assembled into contigs via Megahit v.1.2.9^84^ with ‘meta-sensitive’ parameters, and Prodigal v.2.6.3 on the metagenome mode (−p meta) was then used to predict genes. Subsequently, we utilized CoverM V.4.0 to estimate gene abundance (-m rpkm) by mapping reads to the non-redundant reference constructed with CD-HIT using a sequence identity cutoff of 0.95, and a minimum coverage cutoff of 0.9 for the shorter sequences.

### Estimating the abundance of secondary bile acid production species and genes

Owing to the presence of multi-copy SBA production genes in certain genomes, to mitigate the bias caused by the multiple copies, we weighted the abundance of SBA-production species by the average copy number (Supplementary Table 7) of the genomes within each species(equation (1)(2)). And we compared the weighted relative abundance of these SBA-production microorganisms between diseases and control using two-tailed Mann-Whitney U-test (UC, CD, adenoma, CRC) or paired t test (NAFLD) to find out the differential SBA-production microorganisms.

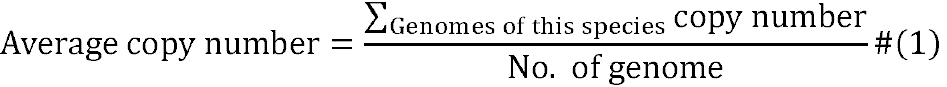

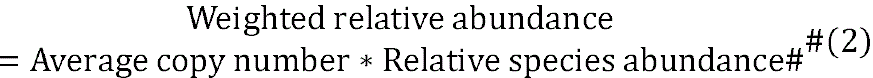

Based on BLASTX searches of genes in the non-redundant reference from metagenomic data against the non-redundant protein catalog of SBA-production, we classified genes from metagenomes as SBA production genes if they exhibited at least 70% identity and an e-value of less than 1e^−50^. The abundance of a certain type of gene was estimated by the sum of the reads per kilobase per million mapped reads (RPKM) calculated by CoverM. Considering the entire metabolic process, we calculated the proportion of Bai genes and HSDHs genes by considering their abundances as a whole. The abundance of BSH and proportion of Bai genes or HSDHs between diseases and control were compared using two-tailed Mann-Whitney U-test (UC, CD, adenoma, CRC) or paired t test (NAFLD) to calculate the p value for determining the significance. Subsequently, combining the importance of various genes and the significance level of their differences (SL in equation (3)), we defined a difference score (equation (3)) to assess the overall dysregulation of BA metabolism from a microbial perspective in diverse diseases.

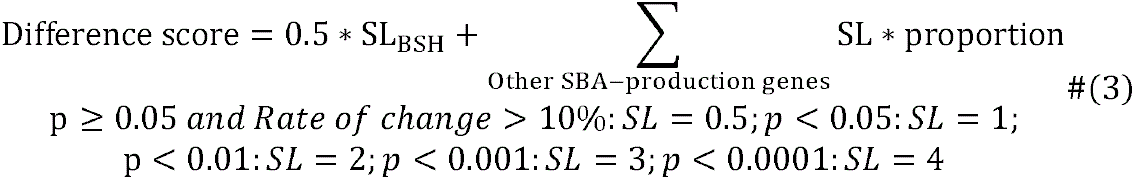

### Comparing secondary bile acid production genes in humans and other animal models

To compare microbial SBA metabolism, the DNA sequences of GMGC unigenes potential for SBA metabolism in the gut of humans and other animal models were compared using BLASTN. For SBA-production unigenes in human, those shared with other animal models were categorized as ‘Same’; for the remaining unigenes, if they had hits with at least 70% identity and less than 1e^−50^ e-value, they were labeled as ‘Similar’, while the rest were classified as ‘Unique’. Then, with the proportions of these three gene categories, scores representing the similarity of specific types of SBA metabolic genes between animal models and humans were estimated using the defined scoring rules based on the match quality (equation (4)). By further assigning different weights on these scores to reflect the contribution of each gene type to the overall metabolic process, we calculated the overall similarity score (equation (5)) for a comprehensive assessment of the similarity.

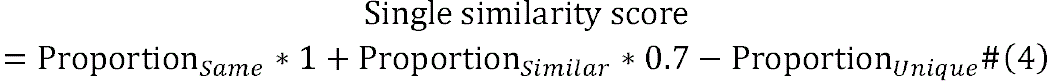

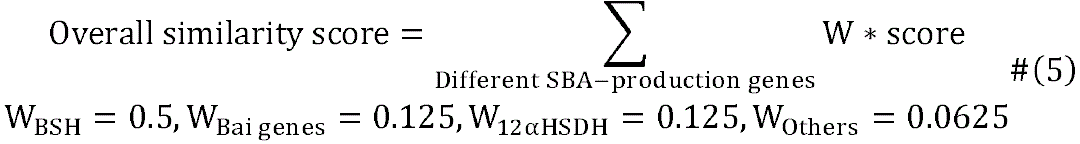

## Data Availability

The comprehensive SBA-production gene catalog of BSHs, Bai genes and HSDHs built based on RefSeq microbiome and its associated data including raw data, the taxonomic and habitat distribution, ect., are available within the paper and its Supplementary Table, as well as on GitHub (https://github.com/ywyang1/SBA-production-gene-catalog/).

Genomes analyzed are available in the RefSeq database (https://ftp.ncbi.nlm.nih.gov/genomes/refseq/, accessed in July 2022). The sequence and metadata of Unigene can be downloaded from GMGC v1.0 (https://gmgc.embl.de/download.cgi). The raw sequencing reads of metagenomic samples were downloaded from SRA of the NCBI database under accession numbers ERP008729, PRJEB1220 and PRJNA420817.

## Code Availability

The scripts for SBA-production gene catalog construction and further analyses are available on GitHub (https://github.com/ywyang1/SBA-production-gene-catalog/).

## Acknowledgements

This work was supported by the National Natural Science Foundation of China (grant number 92251307 to RZ, 82170542 to RZ, 82000536 to NJ), the National Key Research and Development Program of China (grant number 2021YFF0703700/2021YFF0703702 to RZ), and Guangdong Province “Pearl River Talent Plan” Innovation and Entrepreneurship Team Project (grant number 2019ZT08Y464 to LZ). The funders had no role in study design, data collection and analysis, decision to publish, or preparation of the manuscript.

## Author contributions

NJ, RZ and LZ conceived and designed the study. YY and WG drafted the manuscript. RZ, LT, WC, XZ, MS, TX, TZ, XZ, LZ and NJ reviewed and edited the manuscript. All authors read and approved the final manuscript.

## Supplementary information

Supplementary Figure 1

Supplementary Figure 2

Supplementary Figure 3

Supplementary Figure 4

Supplementary Table 1

Supplementary Table 2

Supplementary Table 3

Supplementary Table 4

Supplementary Table 5

Supplementary Table 6

Supplementary Table 7

Supplementary Table 8

## References

1 Radominska, A., Treat, S. & Little, J. Bile acid metabolism and the pathophysiology of cholestasis. Semin Liver Dis 13, 219–234 (1993). 10.1055/s-2007-1007351

2 Kulkarni, M. S., Heidepriem, P. M. & Yielding, K. L. Production by lithocholic acid of DNA strand breaks in L1210 cells. Cancer Res 40, 2666–2669 (1980).

3 Makishima, M. et al. Identification of a nuclear receptor for bile acids. Science 284, 1362–1365 (1999). 10.1126/science.284.5418.1362

4 Xing, C. et al. Roles of bile acids signaling in neuromodulation under physiological and pathological conditions. Cell Biosci 13, 106 (2023). 10.1186/s13578-023-01053-z

5 Alasmael, N., Mohan, R., Meira, L. B., Swales, K. E. & Plant, N. J. Activation of the Farnesoid X-receptor in breast cancer cell lines results in cytotoxicity but not increased migration potential. Cancer Lett 370, 250–259 (2016). 10.1016/j.canlet.2015.10.031

6 Dawson, P. A., Lan, T. & Rao, A. Bile acid transporters. J Lipid Res 50, 2340–2357 (2009). 10.1194/jlr.R900012-JLR200

7 Axelson, M. et al. Bile acid synthesis in cultured human hepatocytes: support for an alternative biosynthetic pathway to cholic acid. Hepatology 31, 1305–1312 (2000). 10.1053/jhep.2000.7877

8 Ridlon, J. M., Kang, D. J. & Hylemon, P. B. Bile salt biotransformations by human intestinal bacteria. J Lipid Res 47, 241–259 (2006). 10.1194/jlr.R500013-JLR200

9 Mallonee, D. H. & Hylemon, P. B. Sequencing and expression of a gene encoding a bile acid transporter from Eubacterium sp. strain VPI 12708. J Bacteriol 178, 7053–7058 (1996). 10.1128/jb.178.24.7053-7058.1996

10 Mallonee, D. H., Adams, J. L. & Hylemon, P. B. The bile acid-inducible baiB gene from Eubacterium sp. strain VPI 12708 encodes a bile acid-coenzyme A ligase. J Bacteriol 174, 2065–2071 (1992). 10.1128/jb.174.7.2065-2071.1992

11 Bhowmik, S. et al. Structural and functional characterization of BaiA, an enzyme involved in secondary bile acid synthesis in human gut microbe. Proteins 82, 216–229 (2014). 10.1002/prot.24353

12 Kang, D. J., Ridlon, J. M., Moore, D. R., 2nd, Barnes, S. & Hylemon, P. B. Clostridium scindens baiCD and baiH genes encode stereo-specific 7alpha/7beta-hydroxy-3-oxo-delta4-cholenoic acid oxidoreductases. Biochim Biophys Acta 1781, 16–25 (2008). 10.1016/j.bbalip.2007.10.008

13 Ridlon, J. M. & Hylemon, P. B. Identification and characterization of two bile acid coenzyme A transferases from Clostridium scindens, a bile acid 7alpha-dehydroxylating intestinal bacterium. J Lipid Res 53, 66–76 (2012). 10.1194/jlr.M020313

14 Bhowmik, S. et al. Structure and functional characterization of a bile acid 7alpha dehydratase BaiE in secondary bile acid synthesis. Proteins 84, 316–331 (2016). 10.1002/prot.24971

15 Funabashi, M. et al. A metabolic pathway for bile acid dehydroxylation by the gut microbiome. Nature 582, 566–570 (2020). 10.1038/s41586-020-2396-4

16 Doden, H. L. & Ridlon, J. M. Microbial Hydroxysteroid Dehydrogenases: From Alpha to Omega. Microorganisms 9 (2021). 10.3390/microorganisms903046917

17 Gadaleta, R. M., Cariello, M., Sabba, C. & Moschetta, A. Tissue-specific actions of FXR in metabolism and cancer. Biochim Biophys Acta 1851, 30–39 (2015). 10.1016/j.bbalip.2014.08.005

18 Zhang, Y. et al. Activation of the nuclear receptor FXR improves hyperglycemia and hyperlipidemia in diabetic mice. Proc Natl Acad Sci U S A 103, 1006–1011 (2006). 10.1073/pnas.0506982103

19 Hang, S. et al. Bile acid metabolites control T(H)17 and T(reg) cell differentiation. Nature 576, 143–148 (2019). 10.1038/s41586-019-1785-z

20 Jiao, N. et al. Suppressed hepatic bile acid signalling despite elevated production of primary and secondary bile acids in NAFLD. Gut 67, 1881–1891 (2018). 10.1136/gutjnl-2017-314307

21 Mohanty, I. et al. The changing metabolic landscape of bile acids – keys to metabolism and immune regulation. Nat Rev Gastroenterol Hepatol (2024). 10.1038/s41575-024-00914-3

22 Yoshimoto, S. et al. Obesity-induced gut microbial metabolite promotes liver cancer through senescence secretome. Nature 499, 97–101 (2013). 10.1038/nature12347

23 Xu, M. et al. Deoxycholic Acid-Induced Gut Dysbiosis Disrupts Bile Acid Enterohepatic Circulation and Promotes Intestinal Inflammation. Dig Dis Sci 66, 568–576 (2021). 10.1007/s10620-020-06208-3

24 Zhong, J. et al. Hyodeoxycholic acid ameliorates nonalcoholic fatty liver disease by inhibiting RAN-mediated PPARalpha nucleus-cytoplasm shuttling. Nat Commun 14, 5451 (2023). 10.1038/s41467-023-41061-8

25 Ruutu, T. et al. Ursodeoxycholic acid for the prevention of hepatic complications in allogeneic stem cell transplantation. Blood 100, 1977–1983 (2002). 10.1182/blood-2001-12-0159

26 Tyagi, P., Edwards, D. R. & Coyne, M. S. Use of Sterol and Bile Acid Biomarkers to Identify Domesticated Animal Sources of Fecal Pollution. Water, Air, and Soil Pollution 187, 263–274 (2007). 10.1007/s11270-007-9514-x

27 Feller, F. M., Holert, J., Yücel, O. & Philipp, B. Degradation of Bile Acids by Soil and Water Bacteria. Microorganisms 9 (2021). 10.3390/microorganisms9081759

28 Philipp, B., Erdbrink, H., Suter, M. J. & Schink, B. Degradation of and sensitivity to cholate in Pseudomonas sp. strain Chol1. Arch Microbiol 185, 192–201 (2006). 10.1007/s00203-006-0085-9

29 Holert, J. et al. Evidence of distinct pathways for bacterial degradation of the steroid compound cholate suggests the potential for metabolic interactions by interspecies cross-feeding. Environ Microbiol 16, 1424–1440 (2014). 10.1111/1462-2920.12407

30 Zhu, Y. G., et al. Ecosystem Microbiome Science. mLife 2, 2–10 (2023). 10.1002/mlf2.12054

31 Faust, K. & Raes, J. Microbial interactions: from networks to models. Nat Rev Microbiol 10, 538–550 (2012). 10.1038/nrmicro2832

32 Heinken, A. et al. Systematic assessment of secondary bile acid metabolism in gut microbes reveals distinct metabolic capabilities in inflammatory bowel disease. Microbiome 7, 75 (2019). 10.1186/s40168-019-0689-3

33 Power, M. E. Top-Down and Bottom-Up Forces in Food Webs: Do Plants Have Primacy. Ecology 73, 733–746 (1992). 10.2307/1940153

34 Kim, G. B., Miyamoto, C. M., Meighen, E. A. & Lee, B. H. Cloning and characterization of the bile salt hydrolase genes (bsh) from Bifidobacterium bifidum strains. Appl Environ Microbiol 70, 5603–5612 (2004). 10.1128/AEM.70.9.5603-5612.2004

35 Wijaya, A. et al. Cloning of the bile salt hydrolase (bsh) gene from Enterococcus faecium FAIR-E 345 and chromosomal location of bsh genes in food enterococci. J Food Prot 67, 2772–2778 (2004). 10.4315/0362-028x-67.12.2772

36 Dussurget, O. et al. Listeria monocytogenes bile salt hydrolase is a PrfA-regulated virulence factor involved in the intestinal and hepatic phases of listeriosis. Mol Microbiol 45, 1095–1106 (2002). 10.1046/j.1365-2958.2002.03080.x

37 Kitahara, M., Takamine, F., Imamura, T. & Benno, Y. Clostridium hiranonis sp. nov., a human intestinal bacterium with bile acid 7alpha-dehydroxylating activity. Int J Syst Evol Microbiol 51, 39–44 (2001). 10.1099/00207713-51-1-39

38 Harris, S. C. et al. Bile acid oxidation by Eggerthella lenta strains C592 and DSM 2243(T). Gut Microbes 9, 523–539 (2018). 10.1080/19490976.2018.1458180

39 Macdonald, I. A., Meier, E. C., Mahony, D. E. & Costain, G. A. 3alpha-, 7alpha– and 12alpha-hydroxysteroid dehydrogenase activities from Clostridium perfringens. Biochim Biophys Acta 450, 142–153 (1976). 10.1016/0005-2760(76)90086-2

40 Coleman, J. P. & Hudson, L. L. Cloning and characterization of a conjugated bile acid hydrolase gene from Clostridium perfringens. Appl Environ Microbiol 61, 2514–2520 (1995). 10.1128/aem.61.7.2514-2520.1995

41 Jones, B. V., Begley, M., Hill, C., Gahan, C. G. & Marchesi, J. R. Functional and comparative metagenomic analysis of bile salt hydrolase activity in the human gut microbiome. Proc Natl Acad Sci U S A 105, 13580–13585 (2008). 10.1073/pnas.0804437105

42 Kim, K. H. et al. Identification and Characterization of Major Bile Acid 7alpha-Dehydroxylating Bacteria in the Human Gut. mSystems 7, e0045522 (2022). 10.1128/msystems.00455-22

43 Song, C., Wang, B., Tan, J., Zhu, L. & Lou, D. Discovery of tauroursodeoxycholic acid biotransformation enzymes from the gut microbiome of black bears using metagenomics. Sci Rep 7, 45495 (2017). 10.1038/srep45495

44 Leviatan, S., Shoer, S., Rothschild, D., Gorodetski, M. & Segal, E. An expanded reference map of the human gut microbiome reveals hundreds of previously unknown species. Nat Commun 13, 3863 (2022). 10.1038/s41467-022-31502-1

45 Guzior, D. V. & Quinn, R. A. Review: microbial transformations of human bile acids. Microbiome 9, 140 (2021). 10.1186/s40168-021-01101-1

46 Coelho, L. P. et al. Towards the biogeography of prokaryotic genes. Nature 601, 252–256 (2021). 10.1038/s41586-021-04233-4

47 Vital, M., Rud, T., Rath, S., Pieper, D. H. & Schluter, D. Diversity of Bacteria Exhibiting Bile Acid-inducible 7alpha-dehydroxylation Genes in the Human Gut. Comput Struct Biotechnol J 17, 1016–1019 (2019). 10.1016/j.csbj.2019.07.012

48 Kim, G. B., Yi, S. H. & Lee, B. H. Purification and characterization of three different types of bile salt hydrolases from Bifidobacterium strains. J Dairy Sci 87, 258–266 (2004). 10.3168/jds.S0022-0302(04)73164-1

49 Lee, J. Y. et al. Contribution of the 7beta-hydroxysteroid dehydrogenase from Ruminococcus gnavus N53 to ursodeoxycholic acid formation in the human colon. J Lipid Res 54, 3062–3069 (2013). 10.1194/jlr.M039834

50 Liu, L., Aigner, A. & Schmid, R. D. Identification, cloning, heterologous expression, and characterization of a NADPH-dependent 7beta-hydroxysteroid dehydrogenase from Collinsella aerofaciens. Appl Microbiol Biotechnol 90, 127–135 (2011). 10.1007/s00253-010-3052-y

51 Doden, H. et al. Metabolism of Oxo-Bile Acids and Characterization of Recombinant 12alpha-Hydroxysteroid Dehydrogenases from Bile Acid 7alpha-Dehydroxylating Human Gut Bacteria. Appl Environ Microbiol 84 (2018). 10.1128/AEM.00235-18

52 Félix, A. P., Souza, C. M. M. & de Oliveira, S. G. Biomarkers of gastrointestinal functionality in dogs: A systematic review and meta-analysis. Animal Feed Science and Technology 283 (2022). 10.1016/j.anifeedsci.2021.115183

53 Correa Lopes, B., et al. Correlation between Peptacetobacter hiranonis, the baiCD Gene, and Secondary Bile Acids in Dogs. Animals (Basel) 14 (2024). 10.3390/ani14020216

54 Clark, V. C. et al. An endogenous bile acid and dietary sucrose from skin secretions of alkaloid-sequestering poison frogs. J Nat Prod 75, 473–478 (2012). 10.1021/np200963r

55 Franzosa, E. A. et al. Gut microbiome structure and metabolic activity in inflammatory bowel disease. Nat Microbiol 4, 293–305 (2019). 10.1038/s41564-018-0306-4

56 Lloyd-Price, J. et al. Multi-omics of the gut microbial ecosystem in inflammatory bowel diseases. Nature 569, 655–662 (2019). 10.1038/s41586-019-1237-9

57 Hogan, S. P. et al. Bacterial Bile Metabolising Gene Abundance in Crohn’s, Ulcerative Colitis and Type 2 Diabetes Metagenomes. PLoS ONE 9 (2014). 10.1371/journal.pone.0115175

58 Eaton, J. E. et al. High-dose ursodeoxycholic acid is associated with the development of colorectal neoplasia in patients with ulcerative colitis and primary sclerosing cholangitis. Am J Gastroenterol 106, 1638–1645 (2011). 10.1038/ajg.2011.156

59 Lindner, S. et al. Altered microbial bile acid metabolism exacerbates T cell-driven inflammation during graft-versus-host disease. Nature Microbiology 9, 614–630 (2024). 10.1038/s41564-024-01617-w

60 Lunney, J. K. et al. Importance of the pig as a human biomedical model. Sci Transl Med 13, eabd5758 (2021). 10.1126/scitranslmed.abd5758

61 Xiao, L. et al. A reference gene catalogue of the pig gut microbiome. Nat Microbiol 1, 16161 (2016). 10.1038/nmicrobiol.2016.161

62 Guilloteau, P., Zabielski, R., Hammon, H. M. & Metges, C. C. Nutritional programming of gastrointestinal tract development. Is the pig a good model for man? Nutr Res Rev 23, 4–22 (2010). 10.1017/S0954422410000077

63 Baker, D. H. Animal models in nutrition research. J Nutr 138, 391–396 (2008). 10.1093/jn/138.2.391

64 Spinelli, V. et al. Influence of Roux-en-Y gastric bypass on plasma bile acid profiles: a comparative study between rats, pigs and humans. Int J Obes (Lond) 40, 1260–1267 (2016). 10.1038/ijo.2016.46

65 Mohanty, I. et al. The underappreciated diversity of bile acid modifications. Cell (2024). 10.1016/j.cell.2024.02.019

66 Guzior, D. V. et al. Bile salt hydrolase acyltransferase activity expands bile acid diversity. Nature 626, 852–858 (2024). 10.1038/s41586-024-07017-8

67 Nie, Q. et al. Gut symbionts alleviate MASH through a secondary bile acid biosynthetic pathway. Cell (2024). 10.1016/j.cell.2024.03.034

68 Krogh, A., Brown, M., Mian, I. S., Sjolander, K. & Haussler, D. Hidden Markov models in computational biology. Applications to protein modeling. J Mol Biol 235, 1501–1531 (1994). 10.1006/jmbi.1994.1104

69 Meng, X. & Ji, Y. Modern Computational Techniques for the HMMER Sequence Analysis. ISRN Bioinform 2013, 252183 (2013). 10.1155/2013/252183

70 Disz, T. et al. Accessing the SEED genome databases via Web services API: tools for programmers. BMC Bioinformatics 11, 319 (2010). 10.1186/1471-2105-11-319

71 Overbeek, R. et al. The subsystems approach to genome annotation and its use in the project to annotate 1000 genomes. Nucleic Acids Res 33, 5691–5702 (2005). 10.1093/nar/gki866

72 Sievers, F. et al. Fast, scalable generation of high-quality protein multiple sequence alignments using Clustal Omega. Mol Syst Biol 7, 539 (2011). 10.1038/msb.2011.75

73 Capella-Gutierrez, S., Silla-Martinez, J. M. & Gabaldon, T. trimAl: a tool for automated alignment trimming in large-scale phylogenetic analyses. Bioinformatics 25, 1972–1973 (2009). 10.1093/bioinformatics/btp348

74 Nguyen, L. T., Schmidt, H. A., von Haeseler, A. & Minh, B. Q. IQ-TREE: a fast and effective stochastic algorithm for estimating maximum-likelihood phylogenies. Mol Biol Evol 32, 268–274 (2015). 10.1093/molbev/msu300

75 Letunic, I. & Bork, P. Interactive Tree Of Life (iTOL) v5: an online tool for phylogenetic tree display and annotation. Nucleic Acids Res 49, W293–W296 (2021). 10.1093/nar/gkab301

76 Camacho, C., et al. BLAST+: architecture and applications. BMC Bioinformatics 10, 421 (2009). 10.1186/1471-2105-10-421

77 Nielsen, H. B. et al. Identification and assembly of genomes and genetic elements in complex metagenomic samples without using reference genomes. Nat Biotechnol 32, 822–828 (2014). 10.1038/nbt.2939

78 Feng, Q. et al. Gut microbiome development along the colorectal adenoma-carcinoma sequence. Nat Commun 6, 6528 (2015). 10.1038/ncomms7528

79 Mardinoglu, A. et al. An Integrated Understanding of the Rapid Metabolic Benefits of a Carbohydrate-Restricted Diet on Hepatic Steatosis in Humans. Cell Metab 27, 559–571 e555 (2018). 10.1016/j.cmet.2018.01.005

80 Bolger, A. M., Lohse, M. & Usadel, B. Trimmomatic: a flexible trimmer for Illumina sequence data. Bioinformatics 30, 2114–2120 (2014). 10.1093/bioinformatics/btu170

81 Langmead, B. & Salzberg, S. L. Fast gapped-read alignment with Bowtie 2. Nat Methods 9, 357–359 (2012). 10.1038/nmeth.1923

82 Wood, D. E., Lu, J. & Langmead, B. Improved metagenomic analysis with Kraken 2. Genome Biol 20, 257 (2019). 10.1186/s13059-019-1891-0

83 Lu, J., Breitwieser, F. P., Thielen, P. & Salzberg, S. L. Bracken: estimating species abundance in metagenomics data. PeerJ Computer Science 3 (2017). 10.7717/peerj-cs.104

84 Li, D., Liu, C. M., Luo, R., Sadakane, K. & Lam, T. W. MEGAHIT: an ultra-fast single-node solution for large and complex metagenomics assembly via succinct de Bruijn graph. Bioinformatics 31, 1674–1676 (2015). 10.1093/bioinformatics/btv033

