## Supplementary Figure 1 for "Systematic identification of secondary bile acid production genes in global microbiome"

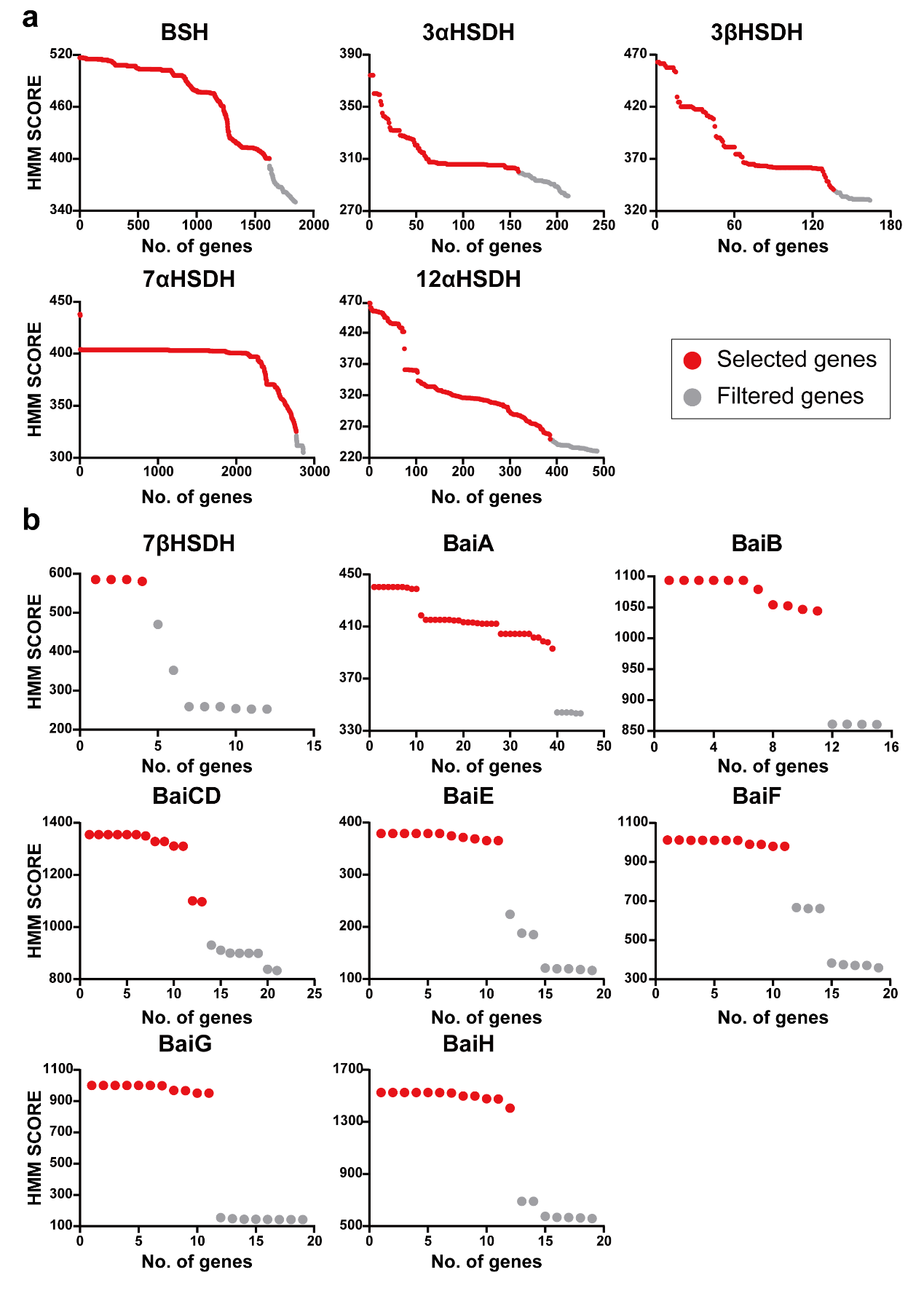

**Supplementary Figure 1. Results of the screening for the secondary bile acid production gene catalog.** (**a**) (**b**) The candidates from hmmsearch are sorted according to their HMM score, and only genes in red are considered as the secondary bile acid production genes. For genes in (**b**), due to limited high-score results, genes from the PubSEED database were included as part of the secondary bile acid production gene catalog.
