## Supplementary Figure 2 for "Systematic identification of secondary bile acid production genes in global microbiome"

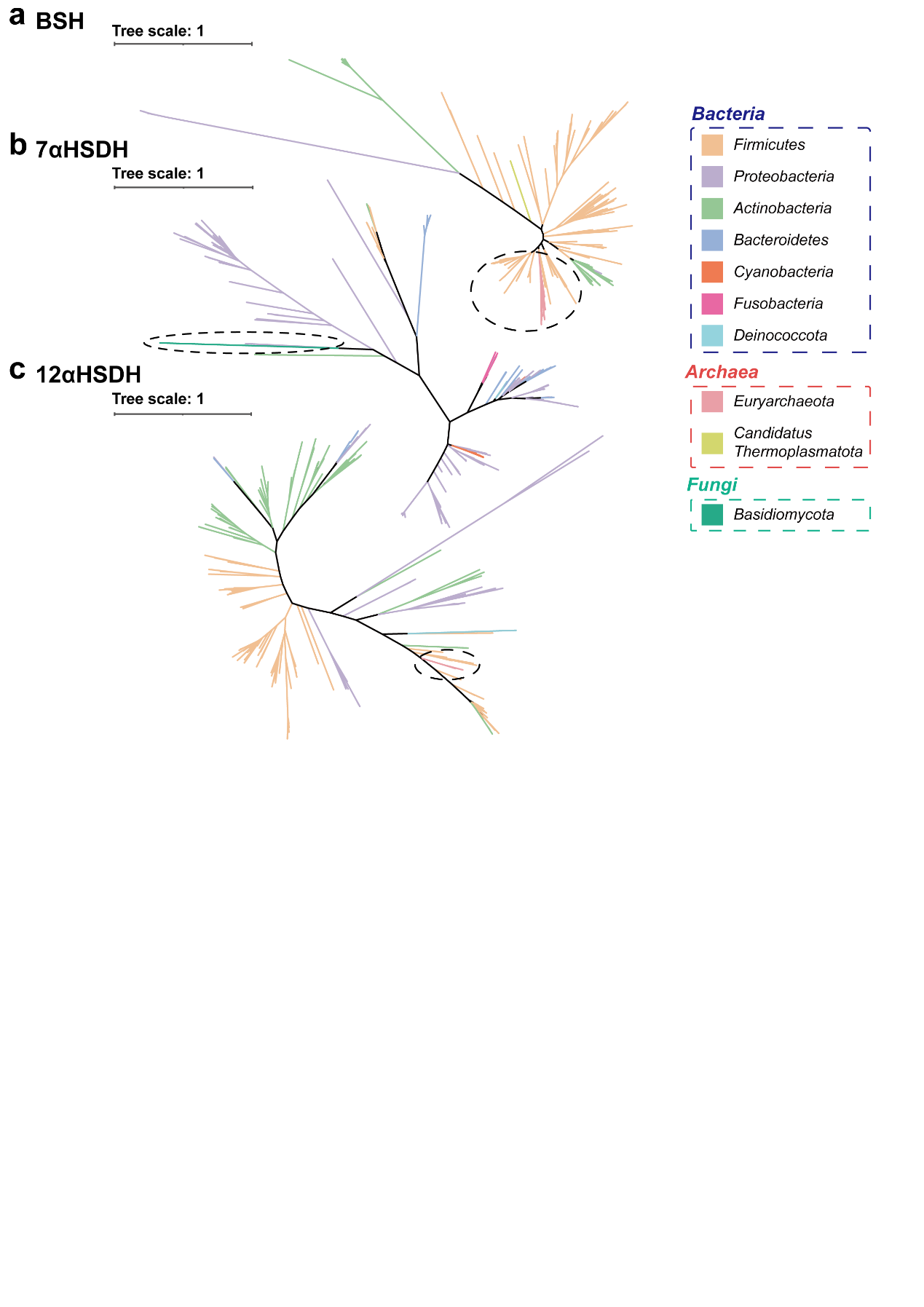

**Supplementary Figure 2. Phylogenetic trees based on the SBA genes distributed in different microbial kingdoms.** Phylogenetic trees based on the non-redundant protein sequences of (**a**) BSH, (**b**) 7αHSDH and (**c**) 12αHSDH. The branch colors represent different phyla.
