## Supplementary Figure 3 for "Systematic identification of secondary bile acid production genes in global microbiome"

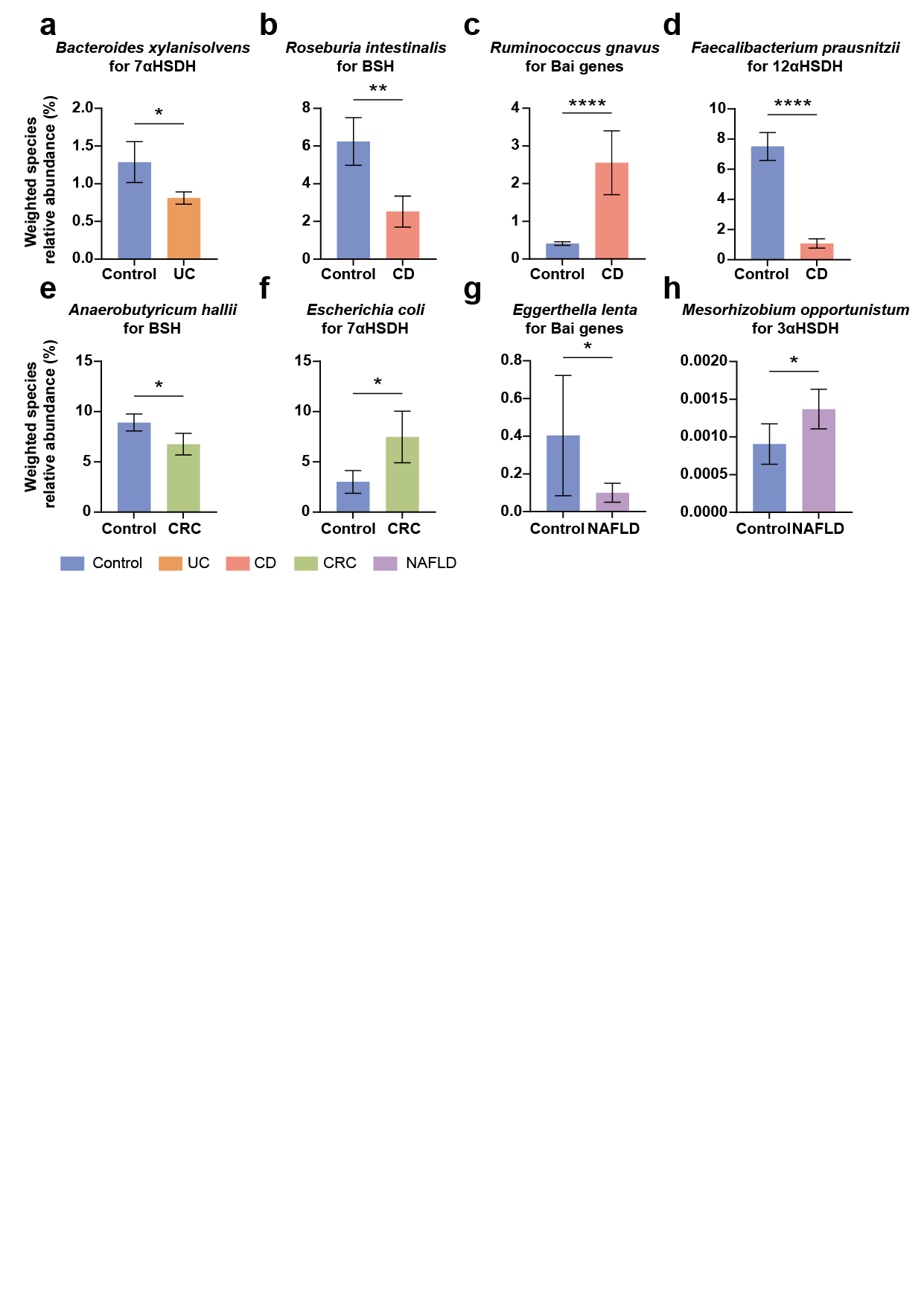

**Supplementary Figure 3. The weighted relative abundance of some major differential species.** The bar plots show the weighted relative abundance of some major differential species. The definition of ‘Weighted relative abundance’ is in equation (2). The bar colors represent different diseases. The definition of ‘Weighted species relative abundance’ is in equation (2). Data are shown as mean with SE. The statistical differences groups were determined by two-tailed Mann-Whitney U-test (UC, CD, adenoma, CRC) or paired t test (NAFLD), the p values were converted to asterisks(* for p ≤ 0.05; ** for p ≤ 0.01; *** for p ≤ 0.001 and **** for p ≤ 0.0001).
