## Supplementary Figure 4 for "Systematic identification of secondary bile acid production genes in global microbiome"

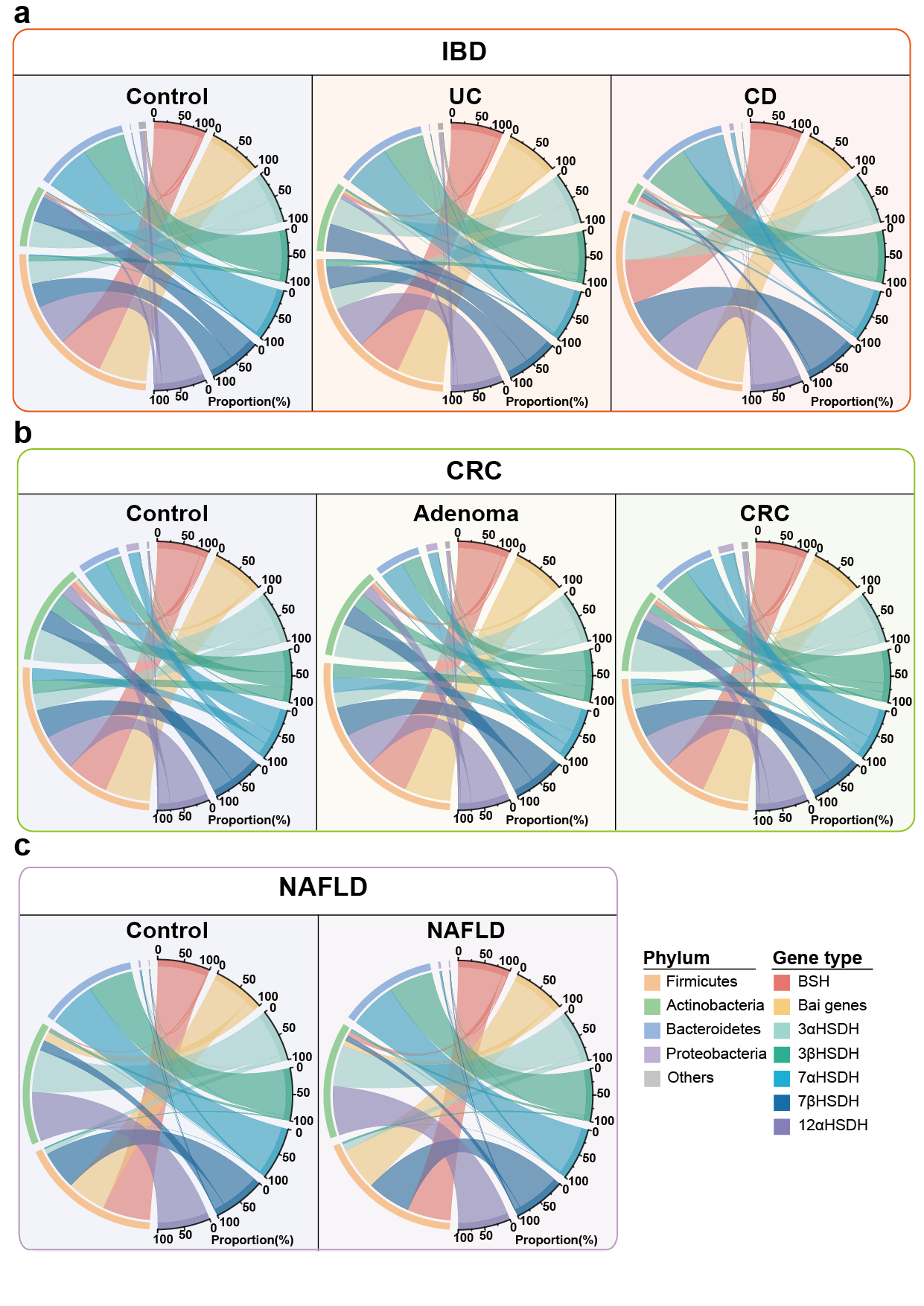

**Supplementary Figure 4. The SBA-production microorganisms composition in intestinal and liver diseases.** The chord diagrams show the weighted species abundance from different phyla related to SBA production. Maximum chord width corresponds to proportion of certain phylum and arrow color designates the gene type.
